# EEG reveals online monitoring mechanisms of speech production

**DOI:** 10.1101/2025.03.01.640939

**Authors:** Joao M Correia, Jill Kries, Lars Hausfeld, Vincent Gracco, Milene Bonte

## Abstract

Speaking involves the orchestration of multiple speech muscles while actively monitoring sensory consequences through auditory and somatosensory feedback. A mistuned sensorimotor mechanism may disrupt the normal integration of motor and auditory brain systems in several developmental and acquired motor speech disorders, including stuttering, speech apraxia, speech sound disorders and dysarthria. Electroencephalography (EEG) provides a non-invasive measure of online neural activity with potential to assess (deficiencies in) sensorimotor integration during speech production. However, the relation between EEG and continuous speechoutput remains poorly characterized. Here, we investigate prediction of auditory speech output using multivariate EEG patterns under three levels of auditory masking. A decoding analysis was employed in combination with a lag-based approach that allowed studying predictions based on instantaneous EEG-speech relations, and their involvement in feedforward and feedback processes. For all masking conditions, we found consistent decoding in instantaneous lags and speech feedback lags, but not in feedforward lags. Furthermore, the level of auditory masking modulated decoding in both the instantaneous and feedback lags. Our results provide insights of neural monitoring during online speech production and offer a window to further study the dysfunction latent in motor speech disorders that may help in optimizing brain-informed therapies for speech fluency.

## Introduction

Speech production is a complex task relying on the orchestration of respiratory, phonatory and articulatory muscles ^1,2^, as well as, the active monitoring of their sensory consequences, including proprioceptive, tactile and auditory feedback ^3^. During production, motor and sensory brain processes are dynamically linked, and unfold continuously and recursively in real-time. This sensorimotor complicity includes a ‘parity’ between articulation and sensory brain systems ^4^, which apart from fluent speech production, support accurate speech perception ^5–7^ and contribute to the phonological skills involved in reading ^8,9^. Feedforward models of speech production ^3,10^ posit the existence of an internal model relating motor and sensory neural codes dynamically ^11^. Learning the sensorimotor relations that build a robust internal model is thus key to normal speech, which may involve exposure to speech sounds from an early age, the integration of speech feedback during babbling ^12^ and the continuous adjustment to anatomical changes across the life-span. Mistuned internal models may in fact relate to disorders of speech, such as persistent developmental stuttering (PDS) ^13^. In PDS, as well as other motor speech disorders (e.g., dysarthria secondary to Parkinson’s disease), improvements are observed when an external rhythm is used to guide action ^14^. In speech, this is often observed in chorus speaking, speech shadowing or metronome pacing ^15,16^. Beyond external rhythms, alterations of auditory feedback ^17^, such as delay or frequency-shift often show immediate but temporary carry-over benefits ^17^. Overall, it is possible that increased neural synchrony to speech rhythm across the sensorimotor brain network is associated with some of the benefits observed. Finding EEG markers of sensorimotor function during fluent speech may in turn stimulate the development of brain-informed therapies for a range of dysfluent motor speech disorders. At present, however, the limited knowledge base needed to study the relation between EEG and continuous speech measures, restricts the ability to characterize and assign EEG synchronization to one’s own speech.

Natural speech unfolds at the syllabic rate, typically below 8Hz (theta rhythm), which is visible in the envelope signal of speech acoustics. During speech perception, neural synchrony to speech envelope permits investigating online speech segmentation and underlying processing mechanisms ^18–20^. Furthermore, recent advances in EEG analysis provide the means to investigate neural synchrony to stimuli or events involved in tasks with an underlying continuous nature, such as inherent in speech perception ^21–23^. Crucially, a lag-based analysis involving the application of varying amounts of asynchrony between dynamic EEG and speech signals allows for a characterization of time-resolved decoding capacity (e.g., Horton et al., 2013). In speech production research, the application of these multivariate decoding methods in respect to the envelope of speech output is promising to uncover feedforward and feedback processes involved in the neural internal model of speech ^11^. For example, a recent electrocorticography (ECoG) study of fluent speakers found a double dissociation between neural synchrony in pre-frontal and superior temporal cortical sites related to feedforward and feedback production mechanisms, respectively ^25^, and a recent extracranial EEG study unraveled conversation-based EEG synchrony between speakers and listeners ^23^. Importantly, multivariate decoding techniques of speech envelope signals provide specific advantages to explore the EEG brain dynamics of continuous speech production: 1) multivariate modelling based on spatial patterns of EEG activity provide increased model sensitivity; 2) slow oscillations avoid large contaminations from muscle artifacts involved in speaking ^26^; 3) after model training, ongoing model validation is performed virtually in real-time, allowing for the fast feedback necessary in ecological conversational conditions; 4) model training can be subject- and session-specific, which is crucial to address the variability seen in speech disorders, as well, as their recovery evolutions.

Here, we investigate synchronization of ongoing EEG to recordings of speech output during online production. Twenty-one participants performed an overt speaking task comprising multiple pronunciations of a set of six English sentences, while EEG and speech output were recorded (Figure 1A). Simultaneously with production, auditory masks (cafeteria noise) parametrically varying in loudness level were presented via headphones to investigate modulation of re-afferent auditory quality. Sentences were presented as text, but production was delayed and based on memory to avoid contamination from reading-related mechanisms that were not of interest. Our analysis scheme involved the computation of a decoding model based on multivariate regression of spatial EEG patterns with respect to the envelope of the corresponding speech recordings (Figure 2). The lag-based approach comprised repeating the decoding procedure through multiple lags of temporal asynchrony (from -500 to +500 ms) between the speech envelope and the EEG signals, producing a temporal profile of envelope reconstruction performance. In order to assure invariance to sentence type, model validation relied on a sentence generalization scheme (e.g., different sentences were used for model training and testing). Based on recent ECoG findings and models of sensorimotor mechanisms, we hypothesized that EEG decoding would be possible in time-lags related with feedforward (i.e., negative lags reflecting speech planning and execution) and feedback processes (i.e., positive lags reflecting speech monitoring). Importantly, we hypothesized that feedback, but not feedforward, processes should be modulated by quality of re-afferent speech (i.e. by auditory masking level). We also expected that, due to facial muscle artifacts, decoding in instantaneous lags (i.e., lags around 0 ms) would be possible, but that these relations should not be affected by auditory mask level.

**Figure 1.**
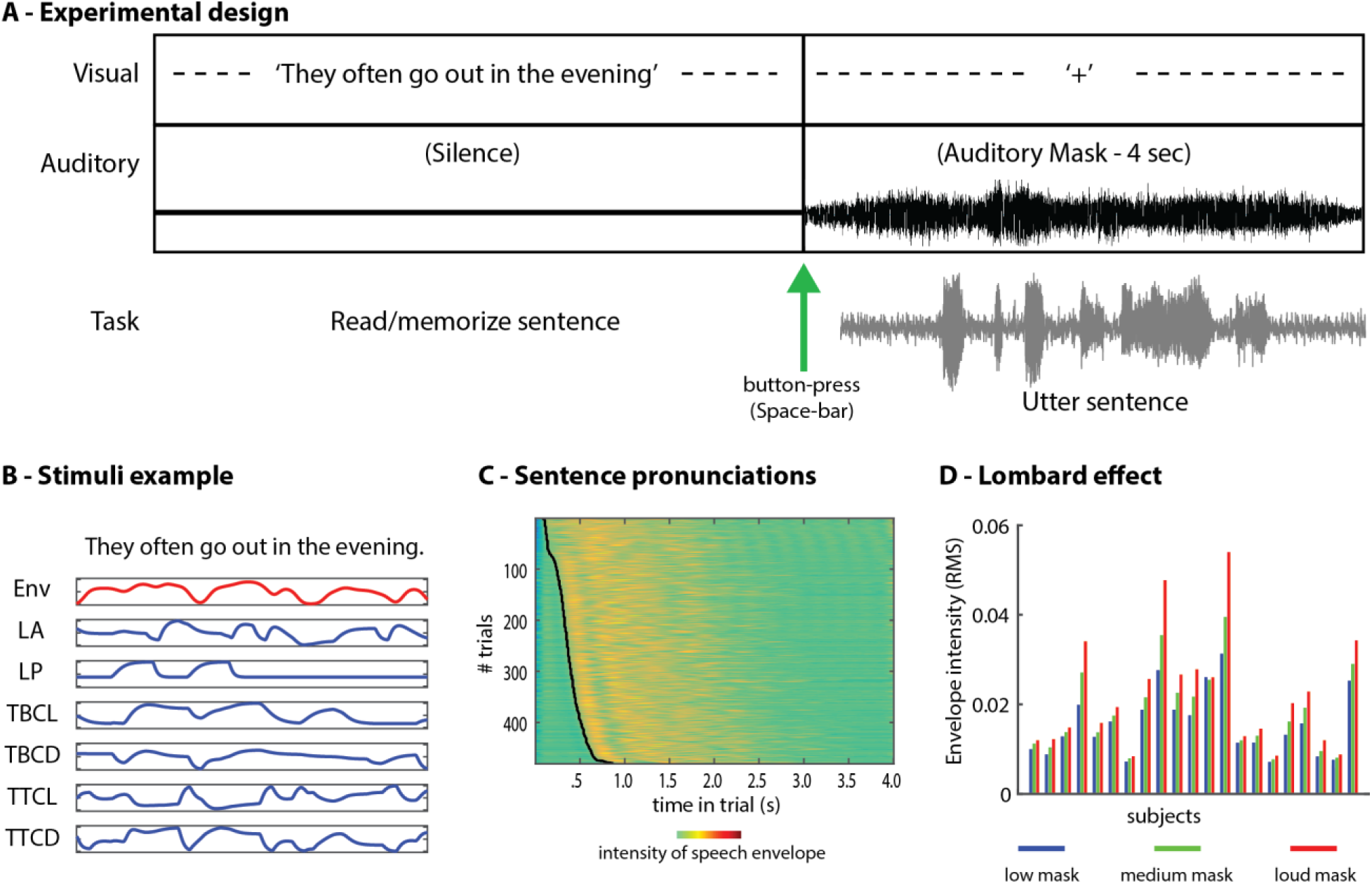
Experimental design and performance. A-Experimental design. During a cueing period, a written sentence was presented in silence. Participants then pressed the space-bar, which replaced the written sentence by a fixation cross and the silence by an auditory mask of cafeteria noise at a random volume level (low, medium or loud). Participants utteredthe sentence from memorywith a fixed limit of 4 seconds, after which the following trial was presented. B-Simulated speech envelope (Env) for a written sentence included in the stimuli, and corresponding articulatory tract variables generated by TADA (TAsk Dynamics Application) software (LA: lip aperture, LP: lip protrusion, TBCL: tongue base constriction location, TBCD: tongue base constriction degree, TTCL: tongue tip constriction location, TTCD: tongue tip constriction degree). C-Stacked envelope signals (z-transformed) uttered across all trials in one of the participants, sortedby speech onset (indicated by black curve). D- Lombard effect observedacross subjects for recordedspeech output within different mask levels. RMS: root mean square.

**Figure 2.**
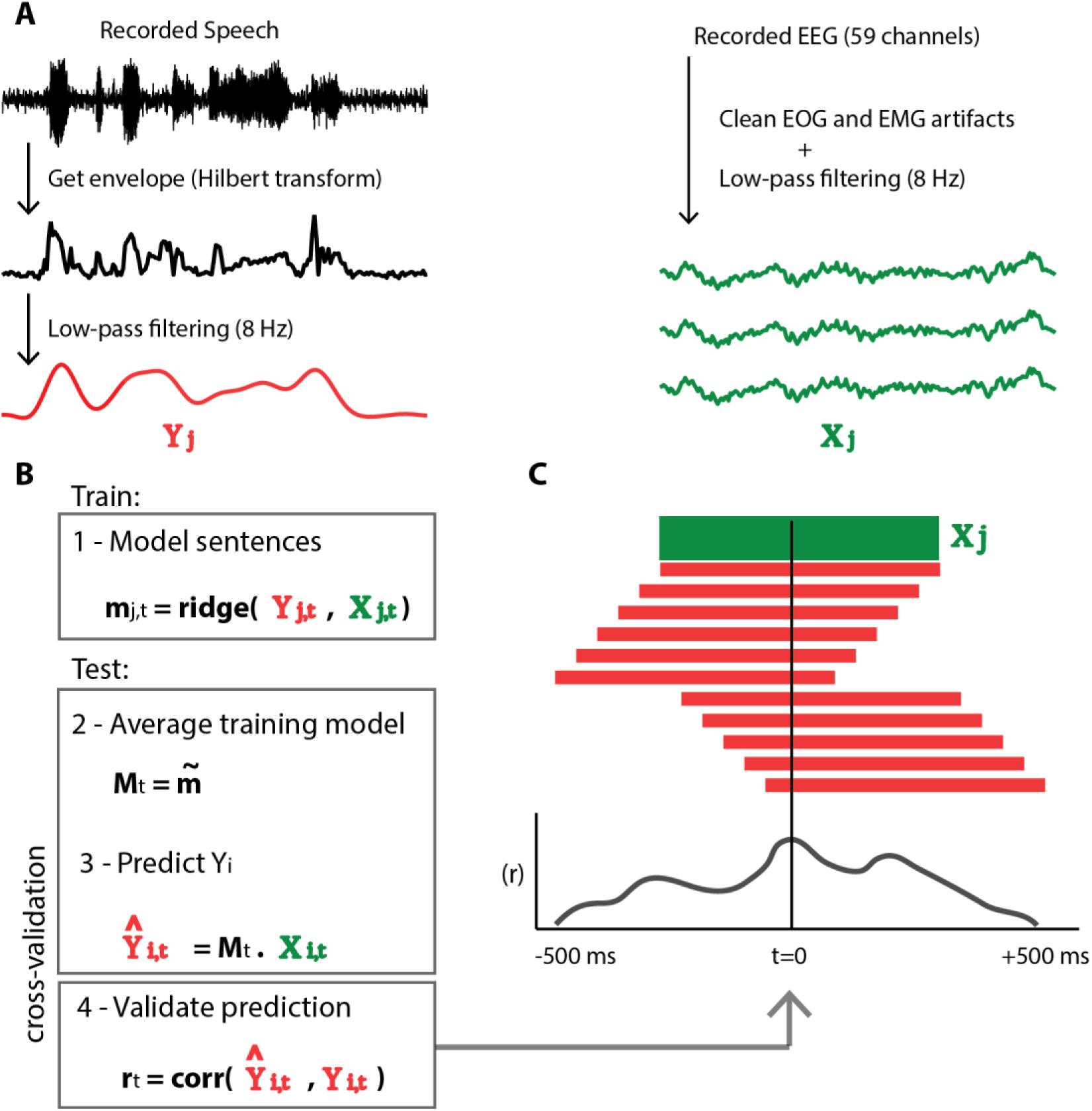
Decoding strategy. A-Computation of the speech envelope (Yj) and preprocessing of the EEG signal (Xj) for a given trial j. B-Decoding analysis was divided into training and testing based on a leave-out-sentence cross-validation procedure. Training consisted on the computation of multiple models (mj,t), for each trial j and lag t, using a spatial pattern of EEG signals across channels (Xj) and the sentence envelope (Yj). Testing consisted of averaging across all training trials for each lag (Mt) followed by prediction of each test trial i for each lag 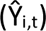. Finally, model performance was assessed using pearson correlation to the original speech envelope of i (Yi,t). C-Averaged decoding performance per subject and cross-validation folds for all lags (-500 to +500 ms). Each lag is determined by the asynchrony of the speechenvelope signal (in red) in relation to the EEG signal (in green). The decoding described in B is repeated for each lag of asynchrony, creating a temporal profile of speech envelope reconstruction. Negative lags represent EEG before speech (or feedforward processes) and positive lags EEG after speech (or feedback processes).

## Methods

### Participants

Twenty-one English speaking right handed adults (10 males, age range 22-43 years, mean 27.95 years, SD 4.54) participated in the study. All participants reported normal hearing abilities and were neurologically healthy. English proficiency was assessed with the LexTALE test (Test for Advanced Learners of English, www.lextale.com ), a vocabulary English test including 40 frequent English words and 20 non-words (Lemhöfer & Broersma, 2012). The mean test score was 86.39% correct (SD=9.01) and all participants were above the average score (70.7%) of a large group of Dutch and Korean advanced learners of English performing the same test (Lemhöfer & Broersma, 2012). The experimental procedures were approved by the ethics committee of the Faculty of Psychology and Neuroscience at Maastricht University (The Netherlands), and were performed in accordance with the approved guidelines and the Declaration of Helsinki. Informed consent was obtained from each participant before conducting the experiments.

### Stimuli

Written stimuli (written sentences) were selected from a large corpus of English written and spoken items (TIMIT Acoustic-Phonetic Continuous Speech Corpus, ^27^). Initially a set of 20 sentences was chosen with comparable number of words, ease of articulation, semantic complexity and emotional load (neutral). Nine independent participants were asked to pilot our task behaviorally using this set of sentences, which helped trimming the final selection to 6 sentences for which no speech production errors were detected in any of the participants. The selected sentences were: “Help celebrate your brother’s success”; “Guess the question from the answer”; “They often go out in the evening”; “We’re open every Monday evening”; “Don’t plan meals that are too complicated”; “Go change your shoes before you turn around”.

Auditory stimuli (auditory masks) consisted of three four-second segments comprising cafeteria noise, stereo recorded with a sampling rate of 44.1 KHz. For each masking segment, 3 distinct versions were created by changing their relative volume using the sound attenuation functionality available in NBS Presentation software (www.neurobs.com) with a ratio of 0.2 for low, 0.1 for medium and 0 for loud noise. The loudest mask was within comfortable level for all the pilot and EEG study participants, but would mask most of their own speech output, whereas the fainter mask would only partially mask their own speech.

### Experimental procedure

The experiment was divided in runs of 60 trials each. Thirteen participants performed 5 runs and eight participants 8 runs. Within each run, each of the 6 sentences was presented 10 times in a pseudo-random order, avoiding consecutive repetitions of sentence or mask level. The experimental design aimed at separating the speech production and active reading phases of the task. Hence, the task consisted of: 1) participants covertly read a sentence that was presented in silence and kept it in memory (no time limit was given for this step); 2) when ready, participants pressed the ‘space-bar’ on the keyboard, which replaced the written sentence by a fixation-cross; 3) the auditory mask was presented together with a visual fixation-cross during a period of 4 seconds; 4) during this period, participants overtly uttered the sentence, which was recorded using a microphone. Four seconds was always sufficient for uttering the sentences, both in the pilot test and the EEG experiment (Figure 1C). Participants were highly familiarized with all the sentences.

### Data recording and pre-processing

EEG data was recorded with a sampling rate of 500 Hz in an electrically shielded and sound proof room from 61 electrode positions (Easycap, Montage Number 10, 10–20 system) relative to a left mastoid reference signal. The ground electrode was placed on the Fz electrode and impedance levels were reduced to minimal levels (below 5 kΩ). EEG preprocessing was performed using EEGlab (Delorme and Makeig, 2004), and included: 1) signal resampling to 250 Hz, band-pass filtering between 1 and 70 Hz and line-noise notch filtering between 48 and 52 Hz; 2) correction of EOG and EMG artifacts using a blind source separation approach based on canonical correlational analysis (or BSS-CCA). BSS-CCA assumes uncorrelated artifact sources, while being maximally correlated to the EEG data, which provides the means to rank decomposed sources based on their autocorrelation and removing them from the data (BSS-CCA toolbox for Matlab, ^28^); 3) filtering the resulting signal in four frequency bands of interest, theta-band limit (1-8 Hz), alpha-band limit (1-13 Hz), beta-band limit (1-30 Hz), and beta-band only (13-30 Hz); 4) removal of EOG channels, re-referencing to the average of left and right mastoid channels and resampling to 100 Hz; 5) epoching of EEG data in relation to onset and offset of speech production identified per trial, followed by baseline correction based on the averaged EEG signal between -100 ms to -50 ms prior speech onset.

Speech utterances were recorded simultaneously with EEG during the 4 second production period of each trial at a sampling rate of 16000 Hz using a single channel microphone. Preprocessing included speech envelope computation and was based on: 1) Hilbert transformation followed by the transformation of all negative values to positive values; 2) resampling to 100 Hz; 3) low-pass filtering using an 8 Hz cut-off.

To avoid modelling the speech envelope based on onset-offset effects, and instead focus on their natural rhythm within the sentences, silent periods occurring before and after each sentence production were removed. Speech onset and offset were identified using a custom-made Matlab routine and later inspected and confirmed visually. The routine followed the following procedure: 1) the contrast of the speech envelope signal was enhanced such that 1% of the signal was saturated at high and low intensities using the matlab routine ‘imadjust’; 2) the signal was binarized using a global threshold value and a normalization value of 0.5 (Otsu, N., "A Threshold Selection Method from Gray-Level Histograms," *IEEE Transactions on Systems, Man, and Cybernetics*, Vol. 9, No. 1, 1979, pp. 62-66.); 3) from the binarized speech envelope signal, small clusters were removed using the matlab routine ‘bwareaopen’ with a minimum region separation of 10 consecutive time points (100 ms); 4) the first and last time points were identified as speech onset and offset, respectively. Trials where speech onset occurred earlier than 100 ms after trial onset or where the duration of uttering was shorter than 1000 ms were rejected from further analysis. The onset was further shifted 200 ms in time to avoid the inclusion of possible speech-onset time-locking effects.

### Lag-based analysis

The lag-based analysis included two parts: 1) modelling the relation between the multi-channel EEG signal and the envelope of the speech signal, which was done using a multiple regression model (Ridge regression) and a cross-validation scheme based on generalization of sentence type; and 2) repetition of this modelling approach over multiple asynchrony lags of the speech envelope in respect to the EEG signal. Ridge regression was conducted for each trial using regularization (λ = 1e^-3^) estimated from optimization performed on the first subject. The goal of regression (Figure 2B) was to predict the speech envelope (Y) using the EEG signal (X) and the model weights (M) for a set of testing trials. Correlation (r) between predicted (Ŷ) and real (Y) speech envelopes was used to assess model performance.

Validation of the decoding procedure relied on cross-validation, which encompassed training and testing in different subsets of the data. Because multiple pronunciations within the same sentence type may show consistency, we adopted a cross-validation scheme based on the generalization of sentence type. For instance, training was based on all trials of sentence 1-5 while testing was restricted to trials of sentence 6. Modelling was performed independently per auditory mask level to allow comparing model performance across auditory masking conditions. Asynchrony between the speech envelope signals and the EEG signal was used to estimate the time-course of envelope prediction during continuous speaking (Figure 2C). At lag=0, no asynchrony was imposed between both signals, thus modelling for this situation reflects the prediction of instantaneous speech envelopes. For negative lags, the speech envelopes were shifted backwards in respect to the EEG signals, thus modelling reflects the capacity of EEG to predict future speech envelopes. Finally, for positive lags, the speech envelopes were shifted forward in respect to the EEG signals, and modelling reflects the capacity of EEG to predict past speech envelopes , or speech monitoring. When creating asynchrony lags, non-overlapping initial or final segments were removed from either signal. Furthermore, model patterns (M) were projected onto a head model by averaging across cross-validation folds and participants. This allowed computing the spatial topography of the averaged models for specific lags of interest (Figure 3C).

**Figure 3.**
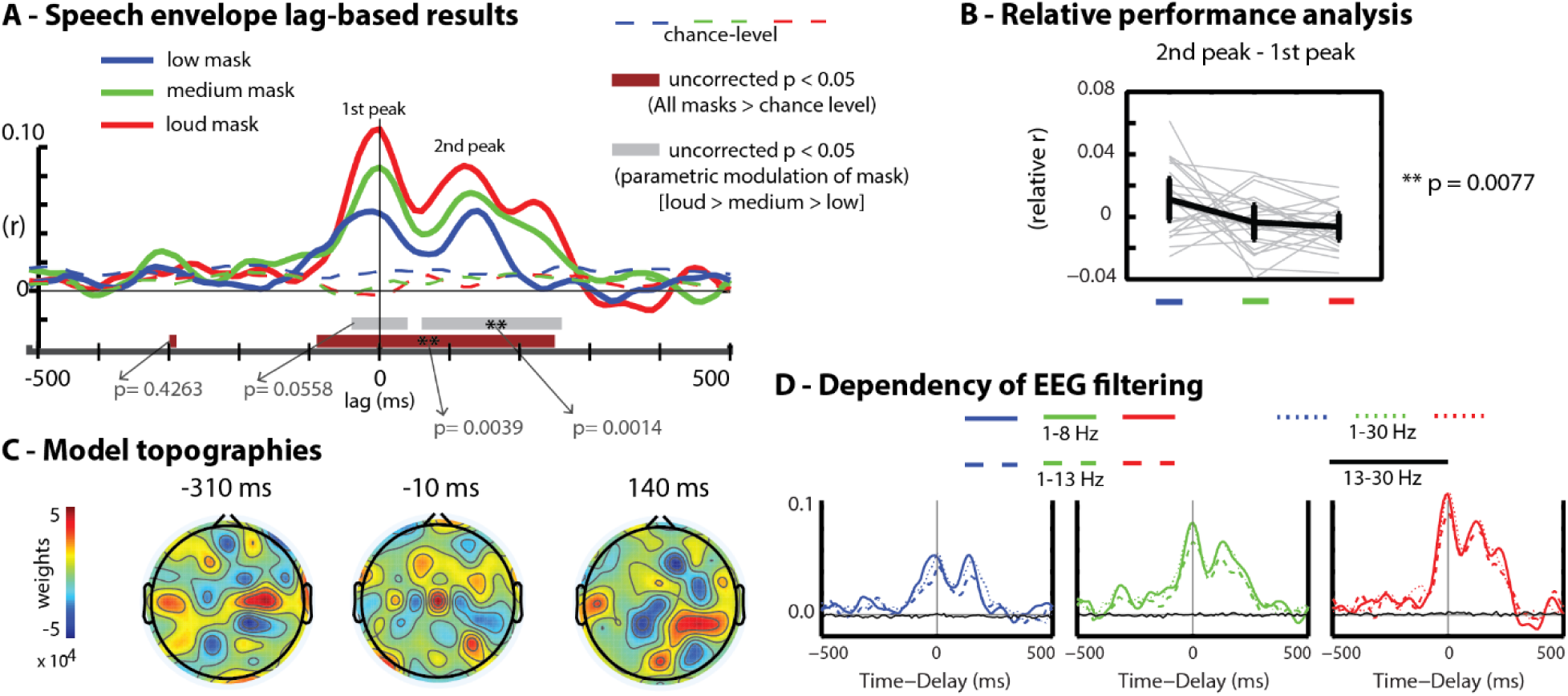
Decoding results of speech envelope. A-Averaged decoding performance of the speechenvelope output from multichannel EEG patterns for each auditory masking volume. Horizontal brown bars indicate overall (all masking levels) significant deviation from permutation chance level (p<0.05, uncorrected). Horizontal gray bar s indicate significant parametric modulations (p<0.05, uncorrected) in respect to masking volume (loud>medium>low). * indicates significant p values after cluster size correction (p<0.05) using a bootstrapping approach (n=10000). B-Relative performance analysis of 2^nd^ peak – 1^st^ peak. Black line represents group average and gray lines individual subject results. C-Averaged topographies of modelling weights for the lag intervals around -310 ms, -10 ms and 140 ms using a window of 100 ms. D-Decoding results dependent on pre-processing filtering bands applied both to the speech envelopes and EEG signals (1 -8 Hz, 1-13 Hz, 1-30 Hz and 13-30 Hz).

### Statistical validation

Individual model performance was computed per subject and lag by averaging across all cross-validation folds. Then, a general linear model (GLM) with three predictors encoding for mask loudness [-1.5 0.5 1] was used to assess the parametric effect of masking volume in our analyses. We also included a GLM with constant predictors [1.0 1.0 1.0] to assess their equal conjoint contribution. Individual empirical chance levels were estimated using cross-trial permutations, that is, for each testing trial, we correlated its predicted speech envelope with the envelope signals of all trials from the other sentence types.

These values were averaged across all possible permutations and cross-validation folds, producing a single correlation value per subject and lag that was subsequently fit to the same GLM predictors described above. Beta values from the real and permuted sets were then used in a second level analysis that tested for group statistical effects using two-sided t-tests. Significant lags (uncorrected p<0.05) were corrected for multiple comparisons (number of lags = 101) based on a maximum cluster size threshold approach using bootstrapping (n=10000) ^29^ and by interchangeably swapping samples between the real and permuted sets (for the model of constant predictors), and by randomly flipping the sign of the beta estimates (for the model of parametric predictors). The distribution of maximum cluster sizes obtained across bootstraps was used to reject the null hypothesis that clusters of uncorrected significant p values occur by chance. We report both uncorrected and corrected p values below 0.05.

### Assessing upper frequency limits

Similar to speech perception studies, our main analysis focused on slow rhythms below 8 Hz that are suitable to track the syllabic rate of speech ^30,31^. In speech production, this frequency band holds the additional benefit of preventing contamination from speech muscle artefacts to the EEG signals ^26^, which occurs mostly in the beta band (13 to 30 Hz). To investigate whether higher frequency limits alter our modelling approach, we varied the low-pass filtering employed in the preprocessing of the EEG and the speech envelope signals, using 8, 13 and 30 Hz. In addition, we applied a band-pass filter (13-30 Hz) to investigate whether beta band oscillations alone could decode the speech envelope signals.

### Decoding of mask sounds and articulatory features of speech

Apart from modelling the speech sound signal, we also investigated the multivariate relation between the EEG signal and the mask sound signal as well as six estimates of articulatory gestures used to utter the spoken sentences (tract-variables). Mask sounds were transformed to sound envelopes and low-pass filtered to 8 Hz using the same method employed for the speech envelope signals, including the same onset/offset limits per trial. Tract variables were estimated using a temporal warping approach ^32,33^ based on a database of English phonemic translations of vocal tract movements. Planning, execution and monitoring of speech production must relate to articulation within the vocal-tract. Since a direct observation of vocal-tract gestures was unattainable in this study, this approach uses instead the acoustics of the recorded speech sounds. Tract variables estimates were lip aperture (LA), lip protrusion (LP), tongue base constriction location (TBCL), tongue base constriction degree (TBCD), tongue tip constriction location (TTCL) and tongue tip constriction degree (TTCD).

## Results

The main analysis of this study focused on modelling the relation between multichannel EEG signals and the envelope of speech sounds acquired simultaneously during speech production. Subjects performed the task fluently, without production errors or delays. Importantly, all subjects showed a Lombard effect. That is, the volume of their speech was increased during louder masks (Figure 1D). The lag-based analysis employed a multiple regression (ridge regression) across asynchrony delays between the envelopes of participants’ speech and simultaneously recorded EEG signals (from -500 to 500 ms). This procedure, in combination with a cross-validation approach based on sentence type generalization and cross-label testing allowed finding temporal trajectories of speech-envelope reconstruction. Figure 3A shows the group average decoding performance for all the lags, separately per auditory mask volume. Statistical analysis indicates lag intervals for which performance was significantly larger than chance level and lags for which reconstruction was parametrically modulated by mask volume (t-tests, p<0.05, uncorrected). Multiple comparison correction was performed using cluster size thresholds obtained from bootstrapping (n=10000) ^29^.

Speech envelope decoding was possible along a lag interval spanning from -100 ms to +250ms (corrected p=0.0039). Within this interval, two prominent peaks are visible (1^st^ peak and 2^nd^ peak). The 1^st^ peak predicting simultaneous speech sounds around 0 ms and the 2^nd^ peak predicting past speech sounds around 140 ms (Figure 3A). Another lag interval was found in a negative lag around -310 ms (uncorrected p<0.05), although not significant after cluster size correction (p=0.4263). Moreover, parametric sensitivity to mask level (i.e., the louder the mask, the better the speech envelope reconstruction) using a GLM fit with predictors [-1.5 0.5 1.0] was observed around 140 ms (corrected p=0.0014) and close to significance around 0 ms lag (corrected p=0.0558). In addition to the direct parametric modulation of mask level, we performed a relative modulation analysis after estimating the performance of the 2^nd^ peak relative to the 1^st^ peak (Figure 3B). This analysis was used to account for the effect observed at 0 ms, which may be influenced by ongoing muscle artifacts. A significant opposite modulatory effect of mask level was then observed, where the lower the mask, the stronger the relative decoding performance at the 2^nd^ peak (p=0.0077).

Furthermore, topographies of regression weights are depicted in Figure 3C using a 100 ms averaging window centered at the above indicated lags of interest (i.e., -310 ms, 0 ms and 140 ms), and averaged across cross-validation folds, subjects and mask volumes. Topographies reflect the weights of the regression models, which indicate the relative importance of the EEG channels for speech envelope decoding at those lags. Because model topographies depict a relative and not an absolute contribution, they should be distinguished from topographies present in more traditional ERP analysis. Nevertheless, they serve as spatial EEG signatures that may be compared to similar decoding studies ^22^.

Frequency band variation was further employed to assess how inclusion of higher frequencies changed the pattern of results. For this purpose, we varied the upper frequency limit to 8, 13 and 30 Hz, and in addition created a filtering version capturing only beta-band components (13-30 Hz). Synchronization can occur at different rhythms of speech, for instance, at the word, syllabic or phonemic level. We found that decoding performance is not affected by the inclusion of higher frequencies above 8 Hz. However, removing oscillations below 13 Hz affected decoding performance at all time lags.

Furthermore, we repeated the decoding analysis using the envelope of the auditory masks (Figure 4B) to investigate synchronization to the fluctuations imposed by auditory stimulation out of interest. Although less prominent than speech envelope decoding, mask envelope decoding was significant within similar lag intervals (i.e., -20 to 250 ms) with a parametric modulation of mask level around 300 ms lags (correct p<0.05). Finally, the same procedure was used to investigate synchronization of EEG signals to estimated articulatory speech gestures (or tract-variables) (Figure 4C-H). The tract-variables tested were lip aperture (LA), lip protrusion (LP), tongue base constriction location (TBCL), tongue base constriction degree (TBCD), tongue tip constriction location (TTCL) and tongue tip constriction degree (TTCD). Each tract variable is an articulatory estimate as a function of time and refers to movement of the respective articulators throughout each production trial (see Figure 1B for an example). Tract-variables and speech envelopes covaried significantly (Figure 4A). Cross-correlation between envelopes, masks and the six tract-variables was computed using Pearson correlations, z-transformed, averaged across trials and statistically assessed using random effect statistics (one-sample t-tests). Decoding of tract-variables using the lag-based approach revealed statistically significant performances. We found lag intervals with significant decoding for most of the tract-variables, especially during simultaneous and positive time lags (i.e., from -100 to 400 ms). For some tract variables, including LA, TBCL, and TTCD, decoding results were similar to those obtained in envelope decoding, reflecting their signal covariance (Figure 4A). In contrast, lip protrusion (LP) showed significant decoding (corrected p<0.05) around 300 ms, but not before. Curiously, tongue-base displacement (TBCL) showed decoding around 300 ms for both louder masking conditions, but not for the low masking condition, as assessed using false discovery rate (FDR) correction (q<0.05).

**Figure 4.**
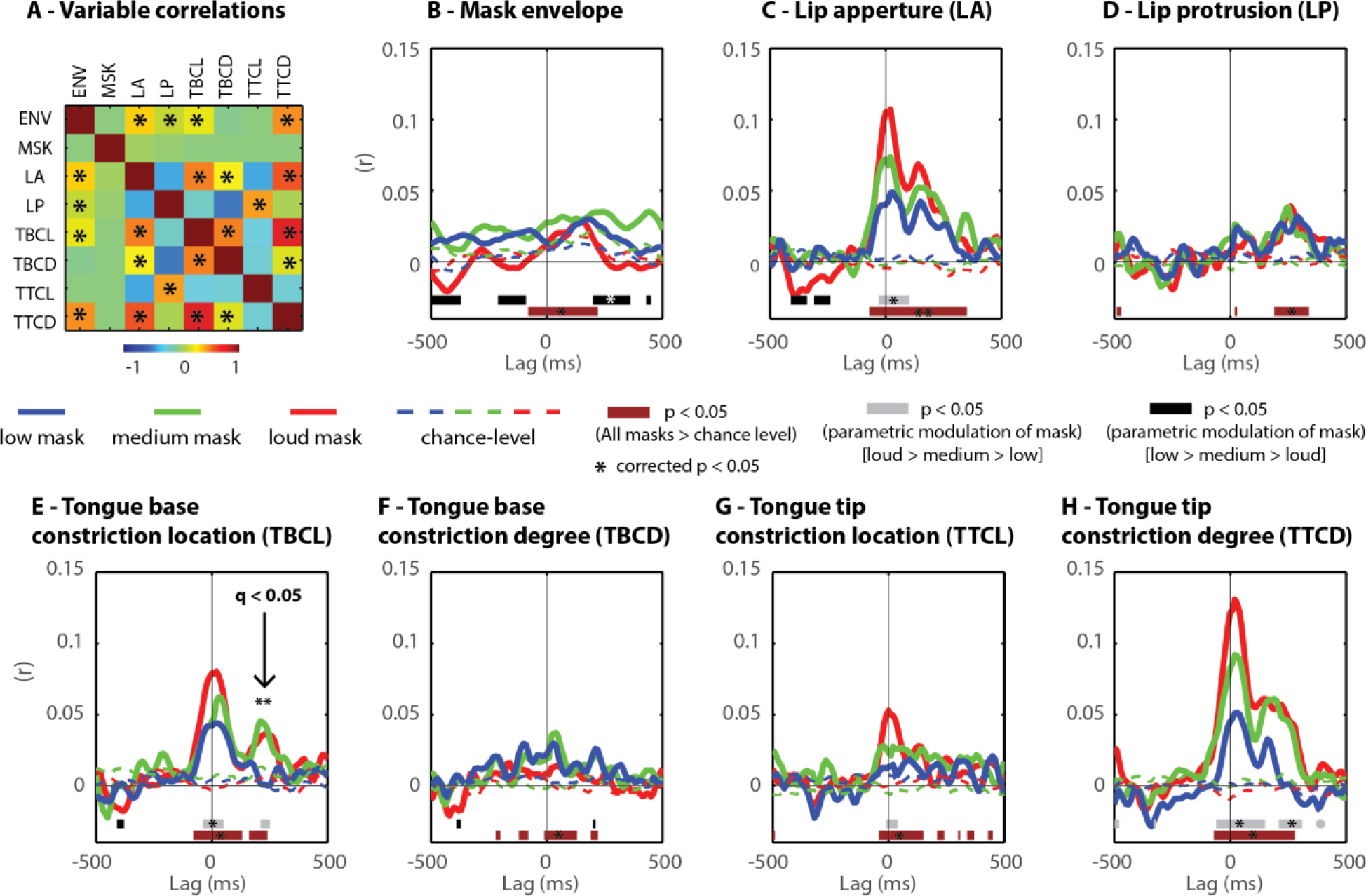
Decoding mask sounds and articulatory features of speech. A- Correlation matrix between the signals of speech envelope (ENV), mask envelope (MSK), as well as the articulatory features tested (LA-lip aperture; LP-lip protrusion; TBCL-tongue base constriction location; TBCD-tongue base constriction degree; TTCL-tongue tip constriction location; TTCD-tongue tip constriction degree). In A, * indicates significant (p<0.05) correlations between signals. B - Decoding the envelope of the auditory masks (MSK). C to H - Decoding of estimated articulatory features. Gray significance bars: loud mask > medium mask > low mask. Black significance bars: low mask > medium mask > loud mask. In B to H, * indicates significant p values after cluster size correction (p<0.05) using a bootstrapping approach (n=10000). Arrow indicates significantly better performance for louder masks in comparison to low mask (q<0.05, FDR).

## Discussion

Investigating the brain dynamics during ongoing speech production is critical to understand how humans master verbal communication, and help uncovering markers of speech fluency that may be used to optimize brain-informed therapies for speech disorders. Brain processes related to the planning, execution and monitoring of speech gestures unfold in parallel during online speech. A brain network based on sensorimotor principles is hence required to integrate these seemingly independent processes, and has been suggested to fail during dysfluent events, such as those observed in individuals who stutter ^13^. In this study, we track these sensorimotor processes by examining the relation between simultaneously recorded EEG and continuous speech output. We used multivariate decoding in combination with a lag-based approach reflecting the time-resolved relation between EEG signals and envelope signals obtained from the participants’ speech. Importantly, our results show significant speech envelope reconstruction at positive time-lags related with speech monitoring, as well as, simultaneous time-lags (Figure 3A). Monitoring of one’s own speech during online conversation relates to the processing of re-afferent speech sound, a critical component of the sensorimotor network of speech production ^3^.

Recently, a univariate correlational analysis between posterior inferior frontal and superior temporal ECoG recordings during overt sentence production distinguished speech planning and monitoring, respectively ^25^. Beyond invasive cortical recordings, multivariate decoding methods have allowed studying the auditory percept using extracranial EEG and MEG (Aiken & Picton, 2008; Horton, D’Zmura, & Srinivasan, 2013; O’Sullivan et al., 2015). Here, we apply similar decoding methods to study neural mechanisms of online sentence production with non-invasive EEG recordings. We demonstrate the feasibility of EEG to uncover feedback monitoring mechanisms of speech production in the absence of ongoing stimulus processing. Like so, our experimental paradigm avoided online reading by separating experimental events related to visual text processing from those related to overt speaking. Subjects kept sentences in memory and produced them in absence of visual stimulation. This strategy was necessary to exclude eye movement confounds involved in visually scanning the stimuli.

Three levels of auditory masking, presented simultaneously via headphones, were employed to assess interference to re-afferent auditory processing. Our initial parametric statistics showed that the louder the mask volume, the better the envelope reconstruction during both, the instantaneous lags (0 ms) and the speech monitoring lags (140 ms). However, in contrast to a disrupted re-afferent speech processing, this tendency (uncorrected p<0.05, corrected p=0.1519) could reflect an EEG response enhancement related to the observed Lombard effect (Figure 1D). In fact, subjects produced louder speech during the louder masking conditions, which influences the intent analysis underlying auditory masking manipulation. To account for the possible influence of the Lombard effect in the auditory masking analysis, we conducted a second analysis based on the relative performance between the two most prominent peaks found by decoding (i.e., around 0 ms and around 140 ms) (see Figure 3A and 3B). While the second peak (140 ms) may relate to speech monitoring processes, given the time delay necessary for re-afferent speech sound processing, the first peak (0 ms) suggests simultaneous prediction of processing speech sounds, which is unlikely related with actual neural processing. In fact, previous intracranical ECoG analyses ^25^ report a time-lag in conformity with the second peak, but not with the first peak. Hence, the first peak could rely on EEG activity patterns reflecting residual muscle artifacts, which can fluctuate with speech intensity reflected in speech volume. The relative performance analysis (Figure 3B) addresses this issue by assessing the modulation of masking level using the performance observed in the second peak relatively to the performance in the first peak. In turn, the lower the mask volume, the better the speech decoding performance in the second peak (around 140 ms lags). These results suggest that sentence production indeed includes tracking of re-afferent speech, which is directly affected by reduced intelligibility imposed by the louder masks. Accordingly, our method provides the means to uncover re-afferent auditory processes during ongoing conversation, a valuable EEG marker of speech monitoring.

On the other hand, our paradigm did not uncover feedforward planning mechanisms, as previously identified by intracranial recordings ^25^. Moreover, the sentences used were simple sequences repeated multiple times, involving a single respiratory event, and possibly encompassing automatic sensorimotor transformations prior to sentence onset. It is possible that longer production events, such as longer sentences, paragraphs or continuous conversation, would provide the additional statistical power to unravel feedforward speech planning mechanisms ^23^. Synchronization to future speech sounds would reflect either the creation of efference copies of speech or their transfer to sensory brain systems, which is supported by feedforward models of speech production ^3,10,34^. It is suggested that efference copies operate at the level of individual words and syllables in the cortex, and possibly at the level of phonemes in the sub-cortex ^3^. Thus, extracranial EEG would potentially allow uncovering ongoing neural mechanisms of speech production related to the word and syllabic rate of speech (typically below 8 Hz).

Furthermore, muscle artefacts normally contaminating the EEG signals during speech production ^26,35^ may have influenced our modeling results. In our main analysis, both the EEG and speech envelope signals were low-pass filtered at the theta band limit (8 Hz), which conservatively removes muscle artefacts ^35,36^. Crucially for the lag-based analysis, muscle artefacts travel at instantaneous rates and are thus expected to influence decoding at lags around 0 ms (i.e., no asynchrony between the signals). Off-center lags, as the ones observed around 140 ms may more reliably point to cognitive processes related with speech monitoring. Alone, beta-band rhythms (13-30 Hz) failed to model speech envelopes, indicating that decoding relies on slower rhythms, including the theta-band.

Apart from speech envelope decoding, we also investigated ongoing mask signal tracking. While uttering the target sentences, cafeteria background sound was delivered using headphones, with the primary goal to mask own-speech feedback but not speech production. Modelling the relation between EEG and the envelope signals of these sounds was initially expected to provide an indication of sound perception mechanisms in positive lags, but not in negative lags, since perception encompasses transmission and processing delays. As expected, this analysis yielded significant results in positive time-lags related with the processing of masking input. Significant decoding performance was observed in positive time-lags, in line with processing of auditory input in previous studies ^21,24^. Nevertheless, overall performance was low and decoding periods spanned inclusive onto instantaneous and negative lags (Figure 4B). This fact may reflect the low temporal variation present in the masking sounds which possibly allowed for sequential predictions. In the future, masking sounds with increased temporal variation, such as music could be also adopted to further disentangle EEG synchronization to production from perception processes (Hausfeld et al., *submitted*, 2017).

Understanding the neurobiological dynamics of speech production further requires studying the relation between brain signals and speech gestures. In EEG, comfortable methods for online gestural imaging are not available. However, under controlled speaking conditions, such as spoken sentences, speech gestures may be estimated from speech acoustics ^32^. Here, this allowed investigating EEG decoding to articulatory estimates of lip and tongue movements (Figure 4C-H). We found selective EEG modulations at specific lag intervals, as well as, distinct modulatory effects of masking. For instance, we found an interesting effect of mask level for tongue-base-constriction-location (TBCL) around a lag centered at +300 ms. This effect shows that monitoring of tongue-base movements is enhanced for louder masks and almost non-existent for the faintest mask level, reflecting engagement of somatosensory monitoring as a compensatory mechanism when auditory monitoring is compromised. At instantaneous lags, decoding individual tract-variables, related with articulatory muscle control, seem to be related with the spatial coverage of the underlying muscles. For instance, lip-protrusion (LP) involves the muscle orbicularis-oris, encircling the lips, which has no overlap with EEG recording sites ^37^. Conversely, tract-variables (e.g., lip-apperture, tongue-base location and tongue-tip constriction degree) relating or covarying with more posterior speech muscles (e.g., the muscle masseter involved in jaw movements) showed higher decoding at instantaneous lags. Beyond EEG, the spatio-temporal brain dynamics of speech gestures may be studied using magnetoencephalography (MEG) in combination with a recent MEG-EMA (electromagneto articulography) technique ^38^, and the spatial organization of the motor speech system using functional MRI (fMRI) in combination with an fMRI-rtMRI (real-time MRI) technique ^39^. These techniques hold promise to unravel details of the sensorimotor mechanisms required for articulatory control when used conjointly with decoding methods and ongoing speech production paradigms, namely providing differentiation between speech and matched non-verbal articulatory gestures.

To conclude, this study investigated sensorimotor processes of online speech production using EEG, a controlled selection of spoken sentences, and a multivariate decoding analysis previously employed to investigate the speech and auditory percept ^22^. We show that these methods are fruitful to tackle feedback (monitoring) processes of speech production, but not feedforward (planning) processes. EEG is a non-invasive neuroimaging technique, broadly available in research and medical institutes, which show capacity to dynamically interface with computers and guide neurofeedback strategies (Muller et al., 2008). In the future, EEG markers of sensorimotor processes required for fluent speech production may significantly impact our ability to design neurofeedback therapies for motor speech disorders ^41^. Although there are multiple possible ways to obtain and deliver feedback for speech production, including altered auditory feedback and multiple sources of bio-feedback, desired carry-over effects crucial for long-lasting benefits may require change at the level of the neurological underpinnings of the dysfluencies. This includes testing and developing subject-specific markers that are flexible to adjust across training sessions.

## Data Availability

The datasets generated and analysed during the current study are available from the corresponding author on request.

